# GAMYB modulates *bHLH142* and is homeostatically regulated by TDR during rice anther tapetal and pollen development

**DOI:** 10.1101/2020.09.26.314641

**Authors:** Swee-Suak Ko, Min-Jeng Li, Yi-Cheng Ho, Chun-Ping Yu, Ting-Ting Yang, Yi-Jyun Lin, Hung-Chien Hsing, Tien-Kuan Chen, Chung-Min Jhong, Wen-Hsiung Li, Maurice Sun-Ben Ku

## Abstract

GAMYB, UDT1, TIP2/bHLH142, TDR, and EAT1/DTD are important transcription factors (TFs) that play a crucial role during rice pollen development. This study demonstrates that bHLH142 acts downstream of UDT1 and GAMYB and works as a “hub” in these two pollen pathways. We show that GAMYB modulates *bHLH142* expression through specific binding to the MYB motif of *bHLH142* promoter during early stage of pollen development; while TDR acts as a transcriptional repressor of the GAMYB modulation of *bHLH142* by binding to the E-box close to the MYB motif on the promoter. The up- and down-regulation of TFs highlights the importance that a tight, precise, and coordinated regulation among these TFs is essential for normal pollen development. Most notably, this study illustrates the regulatory pathways of GAMYB and UDT1 that rely on *bHLH142* in a direct and an indirect manner, respectively, and function in different tissues with distinct biological functions during pollen development. This study advances our understanding of the molecular mechanisms of rice pollen development.

**Highlight:** GAMYB can directly modulate the transactivation of the *bHLH142*, but the modulation is repressed by TDR to keep the homeostasis of *bHLH142* gene expression to ensure normal pollen development.

## Introduction

Pollen development during reproductive growth is a precise and complex process, affecting crop fertility and productivity. Therefore, gaining knowledge about the regulatory network of pollen development is critical for laying the groundwork for controlling pollen fertility in rice and production of hybrid rice. The tapetum layer is the innermost layer of the anther wall that provides nutrients for the development of microspores (Goldberg *et al*., 1993). Timely control of tapetal program cell death (PCD) is crucial for pollen maturation (Ko *et al*., 2017). Several transcription factors (TFs) are associated with tapetal PCD, including UNDEVELOPED TAPETUM1 (UDT1, bHLH164) (Jung *et al*., 2005), GIBBERELLIN MYB GENE (GAMYB) (Aya *et al*., 2009), TDR INTERACTING PROTEIN2(TIP2, bHLH142) (Fu *et al*., 2014; Ko *et al*., 2014), TAPETUM DEGENERATION RETARDATION (TDR, bHLH5) (Li *et al*., 2006), and ETERNAL TAPETUM 1/DELAYED TAPETUM DEGENERATION (EAT1/ DTD, bHLH141) (Ji *et al*., 2013; Niu *et al*., 2013). GAMYB is involved in Gibberellin-regulated gene expression in anthers (Aya *et al*., 2009). EAT1/DTD regulates the gene expression of two aspartic proteases, *AP25* and *AP37*, to execute tapetal PCD, and *OsC4, OsC6*, and *RTS* to maintain pollen wall development (Niu et al., 2013; Ji et al., 2013). Another TF PERSISTANT TAPETAL CELL1 (PTC1) also controls PCD and pollen wall development in rice (Li *et al*., 2011).

Lipid biosynthesis is important for the formation of pollen cell wall exines and Ubisch bodies in anthers (Huysmans *et al*., 1998). It has been reported that CYP703A3, a cytochrome P450 fatty acid hydroxylase, is an essential enzyme in sporopollenin biosynthesis, and its expression is regulated by GAMYB (Aya *et al*., 2009). CYP703A3 acts downstream of bHLH142 in the regulatory hierarchy (Aya *et al*., 2011; Ko *et al*., 2014). In addition, OsC6 (LTPL68) and OsC4 (LTP44) coding for lipid transfer protein precursors are also involved in the synthesis of pollen exines during anther development (Tsuchiya *et al*., 1994; Zhang *et al*., 2010). Consistently, YYI (LTP45) and Fatty acyl-CoA reductase (MS2), are downregulated in both *ms142* and *ptc1* mutants (Ko *et al*., 2014; Li *et al*., 2011). Rice polyketide synthase family protein PKS1 (YY2) is also known to function in sporopollenin metabolism (Wang *et al*., 2013; Zou *et al*., 2017). These proteins are essential for pollen maturation. During pollen development immature pollen grains are produced from microspores after meiosis. Meiotic recombination is important to exchange genetic information and create genetic diversity (Li *et al*., 2005). RAD51 plays an important role in homologous recombination during meiotic process (Shinohara *et al*., 1992).

It has been reported that GAMYB functions in parallel with UDT1 in rice to regulate early anther development (Liu *et al*., 2010). *TDR* transcript is decreased in *gamyb-2* mutant, indicating that TDR functions downstream of GAMYB (Aya *et al*., 2009; Liu *et al*., 2010). Our previous gene hierarchy analysis suggested that bHLH142 acts downstream of GAMYB and UDT1 but upstream of EAT1 in rice pollen development (Ko *et al*., 2014). In addition, we have also shown that bHLH142 interacts with TDR to form a heterodimer and then co-modulate the expression of *EAT1* expression (Ko *et al*., 2014). Moreover, overexpression of homologous *bHLH142* leads to premature upregulation of *EAT1* transcription and causes early tapetal PCD, suggesting that timely and precise control of bHLH142 level is critical to normal rice pollen development (Ko *et al*., 2017).

Past studies have extended our knowledge of the complex molecular regulatory cascades during pollen development. However, the interactions among the regulatory TFs identified so far and their positions in the regulatory hierarchy remain unclear. The present study extends our understanding of the underlying mechanisms of the interactions between GAMYB, TDR and bHLH142 during rice pollen development. We provide strong evidences from mutagenesis analysis, transient promoter assay (TPA), electrophoresis mobility shift assay (EMSA), and molecular dynamic simulation of TF-promoter interaction to support our findings.

## Materials and methods

### Plant materials and growth conditions

Rice (*Oryza sativa*) seeds of the *ms142* and *udt1* mutants in TNG67 background were obtained from the TRIM library (http://trim.sinica.edu.tw/). Mutants of *gamyb-2* (in Nipponbare background) and *eat1* (in Hitomebore background) were requested from the National Institute for Agrobiological Sciences (NIAS), Japan. Mutant of *tdr* was obtained from POSTECH Pohang, Korea. Rice seeds of mutants and their wild-type were sown in vermiculite for 3 weeks and then transplanted into soil in the Academia Sinica-Biotechnology Center in Southern Taiwan greenhouse for genetically modified organisms, Tainan, Taiwan.

### Rice transformation

Constructs of the GAMYB (Os01g0812000) and TDR (Os02g0120500) full-length cDNA were PCR amplified using primer sets of GAMYB-BamHI-F and GAMYB-Sal-R, TDR-BamHI-F and TDR-SalI-R, respectively (Table S1). The PCR product is a 1688 bp and 1671 bp fragment, respectively. The fragment was digested with BamHI and ligated into pCAMBIA1390 vector containing the maize ubiquitin promoter. Expression of the selection marker *HptII* gene that encodes hygromycin phosphotransferase was driven by cauliflower mosaic virus (CaMV) 35S promoter. All constructs were confirmed by DNA sequencing. The plasmids were separately transformed and selected by antibiotic. *Agrobacterium tumefaciens* strain EHA105 was used to transfect TNG67 calli as described previously (Chan *et al*., 1993).

### Total RNA isolation and qRT-PCR

Total RNA was isolated from rice tissues using TRIzol Plus RNA purification kit (Invitrogen) as described by the supplier. The stages of anthers were classified into the two stages according to spikelet length: meiosis (4 mm) and young microspore (YM, 6 mm). Total RNA was treated with DNase (Promega), then 1 µg RNA was used to synthesize the oligo(dT) primed first-strand cDNA using the M-MLV reverse transcriptase cDNA synthesis kit (Promega). Quantitative RT-PCR was performed using a CFX96 Real-Time PCR detection system (Bio-Rad, USA). Quantification analysis was performed using CFX Manager Software (Bio-Rad, USA). Primers used for qRT-PCR are listed in the Table S1. *Ubiquitin-like 5* (Os01g0328400) was used as an internal control for normalization of gene expression levels. Each sample had three biological repeats.

### Transient promoter assay

Transient promoter assay (TPA) was performed as previously described (Ko *et al*., 2014) with some modifications. In brief, protoplasts were isolated from young leaf tissue of 10-d-old rice seedlings. The reporter plasmid contained the CaMV35S minimal promoter and the *bHLH142* promoter (3 Kb) fused to the firefly *luc*iferase gene (*LUC*). In the effector plasmids, *GAMYB* and *TDR* were under the control of the CaMV35S promoter. The pBI221 vector containing a CaMV35S promoter driving the expression of GUS was used as an internal control. To verify whether TDR might repress GAMYB modulation on the *bHLH142* promoter, different molar ratios of TDR (1x, 5x, and 10x) and GAMYB proteins were used in the TPA.

### Electrophoretic mobility shift assay

Rice GAMYB and TDR proteins were expressed as a Maltose binding protein (MBP) fusion in *E. coli*. A total of 25 µg protein was incubated with the appropriate Dig-labelled DNA probes. Two fragment of the *bHLH142* probes were synthesized. Probe 1 contained a 66-mer oligonucleotide (−59 to -124 bp from the TSS) was synthesized (Table S1). It contains the MYB-like binding motif 5’-CAACAAA-3’, at -103 to -109 bp from the TSS of *bHLH142*, and an E-box on the *bHLH142* promoter at -73 bp to -78 bp from the TSS of *bHLH142*. Probe 2 from -977 to -1034 bp TSS, consisted of GARE and E-box, was synthesized (Table S1).

### Prediction of 3D protein structures of transcription factors and their interaction with promoters by molecular dynamics simulation

The 3D protein structures of TDR, GAMYB and bHLH142 were constructed by MD simulation using *ab initio* modeling methods. MD simulation was performed by the NAMD (Nelson *et al*., 1996) program using parameters adopted from the CHARMM force field (Brooks *et al*., 1983). The full-length protein sequences of GAMYB (BAF06506.1), and TDR (NP_001045710.1) were obtained from the NCBI Information database. The DNA probes for binding to the TF proteins were generated by 3D-DART server (van Dijk and Bonvin, 2009). The models were minimized by removing unfavorable contacts, brought to 310K by velocity rescaling, and equilibrated for 1 ns. Before any MD trajectory was run, 40 ps of energy minimization was performed to relax the conformational and structural tensions. The minimum structure was the starting point for MD simulation. For this purpose, the protein molecule was embedded into a water simulation box and a cutoff distance of 12 Å was employed for the non-bonded and electrostatic interactions. The heating process was performed from 0 to 310K through Langevin damping with a coefficient of 10 ps21. A time step of 2 fs was employed for rescaling the temperature. After 20 ps of heating to 310K, equilibration trajectories of 2 µs were recorded, which provided the data for the structural and thermodynamic evaluations. The equations of motion were integrated with the Shake algorithm with a time step of 1 fs. Fig. 4 displays atomistic pictures of molecules generated using UCSF Chimera (Pettersen *et al*., 2004).

### RNA Fluorescent *in situ* Hybridization (FISH)

For synthesis of *bHLH142* probe, gene specific primer sets of 5’-CATGTTCAACACCAAGA TTCATTCG-3’ and 5’-TGCAAACCATGACATACCAAAGATC-3’ (Table S1), the same probe sequences as in our previous study(Ko *et al*., 2014), were used. We used DIG RNA labelling kit (SP6/T7) (Cat. No. 11175025910, Roche) to synthesize probes according to the manufacturer’s instructions. FISH was performed, according to previous protocols(Jandura *et al*., 2017) with some modifications. Briefly, primary anti-DIG-POD, Fab fragment (Cat. no. 11207733910, Roche), and tyramide signal amplification (TSA) of the TSA Plus Cyanine 5 (Cy5) detection kit (NEL745001KT, PerkinElmer) were used for a single color FISH experiment.

Anthers at the early meiosis stage of TNG67 and homozygous mutant lines (*ms142, gamyb-2*, and *udt1*) were collected, fixed, dehydrated, embedded, tissue section to 10 μm, and performed *in situ* hybridization. Tissue samples were dewaxed, re-hydrated, protease (Cat. no. 03115836001, Roche) digested, glycine treated (UR-GLY001, Sigma-Aldrich), post-fixed, acetylated with acetic anhydride in triethanolamine, and then dehydrated. Slides were prehybridized at 37°C for 1 h, hybridized with a probe at 58°C overnight, blocking was performed, incubated with anti-DIG-POD antibody at 37°C for 2 h, and then washed three times (15 min each). After that, tissues were reacted with TSA working solution containing Cy5 Plus Amplification Reagent (300x dilution) at room temperature for 5 minutes, and washed twice. Afterwards, slides were counterstained with 4’, 6-diamidino-2-phenylindole (DAPI) (Cat. no. D9542, Sigma-Aldrich). Florescence signals were observed using a Zeiss LSM710 confocal microscope equipped with a T-PMT under a Cy5 filter at excitation wavelength of 633 nm and emission wavelength of 670 nm onwards, and the DAPI signal was excited by a UV filter at a wavelength of 405 nm.

### Two-Color FISH (DISH)

The DISH technique(Brend and Holley, 2009) was used to show the co-expression of *GAMYB* and *bHLH142* transcripts, as well as *UDT1* and *bHLH142* transcripts in the same tissue of TNG67 anther at the early meiosis stage. To synthesize the *GAMYB* probe, gene specific primer sets of 5’-CGAGCTGGGAGGAGCAAAGGA-3’ and 5’-TCCATGGCCCCTTCTTCAGC -3’ were used. To synthesize the *UDT1* probe, gene specific primer sets of 5’-GAGGAGGTGAAGGTGGAGGATGA-3’ and 5’-CGCGAAGATGTTG CCGTTGA-3’ were used. DIG RNA labelling Kit and Fluorescein (FITC) RNA Labeling Mix (Cat. no. 11685619910, Roche) were used to synthesize probes according to the manufacturer’s instructions. Probes of *bHLH142* and *GAMYB* were simultaneously hybridized with tissue slices overnight at 58°C. Then, anti-FITC-POD was conjugated with *GAMYB* probe overnight at room temperature, and then tissues were reacted with TSA working solution containing TSA Plus Cyanine 3 (Cy3) (NEL744001KT, PerkinElmer) (at 1:100 dilution) at room temperature for 5 minutes, and then washed twice. Afterwards, tissues ware treated with 3% hydrogen peroxide at room temperature for 10 minutes to inactivate the first peroxidase of anti-FITC-POD. After that, anti-DIG-POD was conjugated with *bHLH142* probe at 37°C for 2 hr. Tissues were then stained with TSA working solution containing Cy5 amplification reagent (at 1:300 dilution) at room temperature for 5 min, and washed twice. For DISH experiment to study co-expression of *bHLH142* and *UDT1*, probes of *bHLH142* and *UDT1* were hybridized with tissue slices overnight at 58°C. Anti-FITC-POD, Fab fragment was conjugated with *UDT1* probe overnight at room temperature. Then anti-DIG-POD was conjugated with *bHLH142* probe at 37°C for 2 hr. We also performed dye-swap experiments, reciprocal staining of TSA Plus Cy3 and Cy5 to confirm the consistency of RNA fluorescent signals. Fluorescent signals were observed under Zeiss LSM710 confocal microscope equipped with a T-PMT, using Cy3 or Cy5 filter at excitation and emission wavelengths of 543 nm/600 nm or 633 nm/670 nm.

## Results

### bHLH142 is a “hub” of the GAMYB- and UDT1-mediated pathways

We studied the expression patterns of *UDT1, GAMYB, TDR, bHLH142*, and *EAT1* in the anthers of the rice cultivar TNG67 at the meiosis and young microspore (YM) stages and also the expression pattern of *bHLH142* in mutants defective in one of these TFs using qRT-PCR. Our results indicated that in anthers, except for *EAT1*, the other four TFs were expressed highly at the meiosis stage. Interestingly, *bHLH142* was downregulated, while *TDR* and *EAT1* were upregulated at the young microspore (YM) stage (Fig. 1A). The up- and down-regulation of these TFs during anthers development indicate that precise control of the expression level of these TFs at different developmental stages is critical to normal rice pollen development. Several studies have reported that bHLH142 is located downstream of GAMYB and UDT1 (Fu *et al*., 2014; Ko *et al*., 2014). In this study, we investigated *bHLH142* expression patterns in the anther of *gamyb-2, udt1*, and *ms142* mutants at the meiosis stage. Our qRT-PCR data confirmed that *ms142* is a null mutant and does not express *bHLH142* mRNA in the anther. However, compared to the wild-type (WT), the expression of *bHLH142* was suppressed down to 0.38-fold and 0.35-fold in *udt1* and *gamyb-2* mutants, respectively (Fig. 1B). These data suggest that when *UDT1* is knocked out, the GAMYB-pathway is only partially functional with a reduced expression level of *bHLH142*. Similarly, when *GAMYB* is knocked out, the UDT1 pathway is still functional and some level of *bHLH142* is retained in the anther of *gamyb-2* (Fig. 1B). Taken together, these data suggest that the gene regulatory hierarchy of bHLH142 is located downstream of GAMYB-and UDT1-dependent regulatory pathways and serves as the “hub” of both pathways during rice pollen development.

**Fig. 1.**
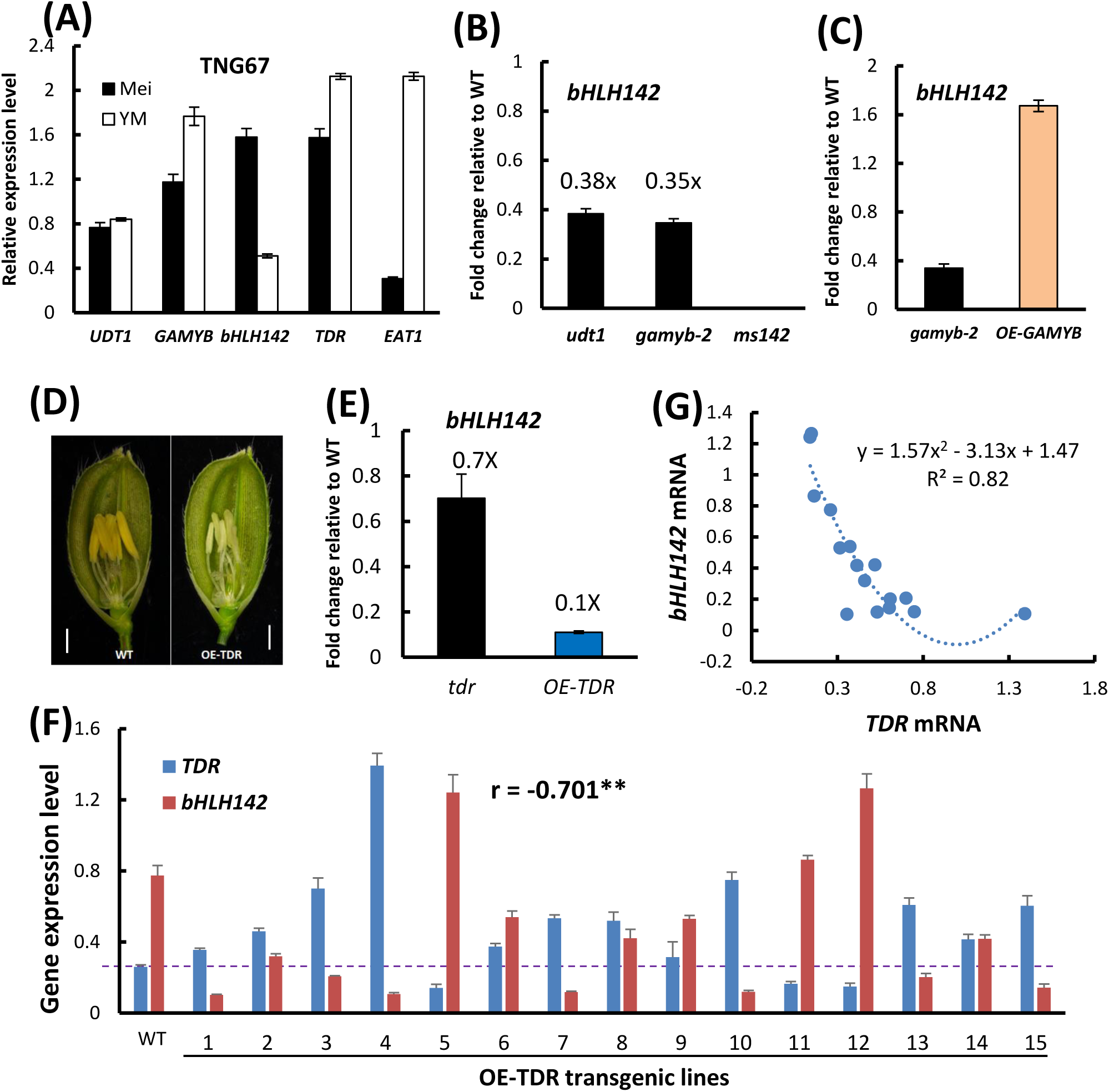
Expression patterns of *bHLH142* and expression relationship between *bHLH142* and *TDR*. (A) Expression patterns of *UDT1, GAMYB, bHLH142, TDR*, and *EAT1* in anthers of TNG67 at the meiosis (Mei) and young microspore (YM) stages. (B) Expression patterns of *bHLH142* in the anthers of *udt1, gamyb-2*, and *ms142* mutants at the meiosis stage. (C) Down- and up-regulation of *bHLH142* in *gamyb-2* and *OE-GAMYB* (line #6) anthers, respectively. (D) Comparison of the spikelet morphology between TNG67 (WT) and OE-*TDR* #4 transgenic rice. The photo was taken one day before anthesis. Bar = 1 mm. (E) Expression patterns of *bHLH142* in the anthers of *tdr* mutant (in Dongjin background) and *OE-TDR* (in TNG67 background) at the meiosis stage. qRT-PCR data are normalized with respective to WT. (F) Expression patterns of *TDR* and *bHLH142* in OE-*TDR* transgenic lines. RNA of anthers at the meiosis stage was collected from 15 independent OE-TDR transgenic lines. (G) Relationship between *bHLH142* and *TDR* mRNA levels in TNG67 (WT) and 15 OE-TDR transgenic lines. Regression analysis fit to a binomial regression equation of y = 1.57x^2^ -3.13x + 1.47 (R^2^ = 0.82); n= 16 lines. Gene expression levels were normalized to *Ubi5* using qRT-PCR. Error bars indicate the SD among three technical replicates. Mei, meiosis; VP, vacuolated pollen. Each experiment was repeated three times.

To test the function of GAMYB, we generated *GAMYB* overexpression lines in the TNG67 background (denoted OE-*GAMYB*) and observed the male sterile phenotype in transgenic rice. OE-*GAMYB* exhibited light yellowish and smaller anthers than the WT. Moreover, pollen grain number was reduced and infertile; consequently, there was no seed developed in the OE-*GAMYB* transgenic lines (Fig. S1). The expression of *bHLH142* was downregulated in the anther of *gamyb-2* mutant, but upregulated in the anther of OE-*GAMYB* transgenic line at the meiosis stage (Fig. 1C). Clearly, loss of function and gain of function in *GAMYB* altered the expression of *bHLH142*.

### Overexpressing *TDR* inhibits *bHLH142* gene expression

We also generated transgenic rice overexpressing *TDR*, as driven by maize *ubiquitin* promoter (denoted OE-*TDR*) for detailed characterization. Similar to OE-*GAMYB*, OE-*TDR* was male sterile and its anthers were small, white yellowish (Fig. 1D), and there was no pollen or seed developed. Our qRT-PCR data indicated that *tdr* knockout mutant had slightly reduced *bHLH142* expression (0.7 x WT) (Fig. 1E). Surprisingly, OE-*TDR* significantly repressed the expression of *bHLH142* (only 0.1 x WT) (Fig. 1E). We further examined the expression levels of *bHLH142* and *TDR* in the anthers of 15 individual OE-*TDR* transgenic lines at the meiosis stage by qRT-PCR and found that in the anther of TNG67, *bHLH142* is expressed at a higher level than *TDR* at the meiosis stage (Fig. 1F). In 12 out of 15 OE-*TDR* lines, *TDR* was upregulated whereas *bHLH142* was downregulated at the meiosis stage, compared to the WT. Except for three OE-*TDR* lines (#5, #11, and #12), transcription of *TDR* was not upregulated, presumably due to the co-suppression effect. It is noteworthy that these three lines showed significant upregulation of *bHLH142* (Fig. 1F) and have normal anther development similar to the WT. Clearly, whenever OE-*TDR* transgenic lines showed an increased *TDR* mRNA, their *bHLH142* transcripts were significantly decreased (r = -0.701**). The expression levels of *TDR* and *bHLH142* were fitted to a negative binomial regression model of y = 1.57×^2^ -3.13x + 1.47 (R^2^ = 0.82) (Fig. 1G). We also assayed the expression patterns of *GAMYB* and *UDT1* in the anther of OE-*TDR* transgenic lines and found no correlation (r=-0.27) (Fig. S2). Using stable transgenic rice overexpressing *TDR*, we demonstrated that the gene expression level of *TDR* is negatively correlated with that of *bHLH142*. Thus, we hypothesize that TDR acts as a repressor in the transactivation of *bHLH142* by GAMYB.

### Identifying DNA-binding motifs of GAMYB and TDR

Since the DNA-binding domains (DBDs) of rice GAMYB and TDR, which belong to the MYB and bHLH families, respectively, have not been determined yet, we searched the TF databases to infer their motifs. The known positional weight matrices (PWMs) were collected from databases, including JASPAR (Khan *et al*., 2018), Cis-BP (Weirauch *et al*., 2014), Plant Cistrome (O’Malley *et al*., 2016), and two papers (Franco-Zorrilla *et al*., 2014; Sullivan *et al*., 2014). The DBD domain sequences of MYB members with known PWMs and the DBD domain sequences of bHLH members with known PWMs in plants were collected. The DBD of GAMYB is most similar to that of AtMYB33 with an 81.6% sequence identity (Fig. 2A), which is higher than the 80% threshold of DBD MYB set by the Cis-BP database, so we assumed that the PWM of GAMYB is similar to that of AtMYB33. The DBD of TDR is most similar to that of AtbHLH13 with a 33.7% sequence identity (Fig. 2A). This sequence identity is lower than the 60% threshold set by Cis-BP, but we also assumed that the PWM of TDR is similar to that of AtbHLH13. We then checked these two putative PWMs by evolutionary conservation and EMSA. For the evolutionary conservation test, we checked the occurrences of the putative motifs of GAMYB and TDR on the 1.5 kb promoter sequence of *bHLH142* from the transcription start site (TSS) in a species using FIMO (Grant *et al*., 2011) with p-value <0.005. We indeed found both putative motifs in the orthologous *bHLH142* promoters in *Leersar perrieri, Sorghum bicolor*, and *Zea mays*, though no putative motif of TDR was found in *Brachypodium distachyon* (Fig. 2B). This evolutionary conservation suggests the functionality of the two putative motifs.

**Fig. 2.**
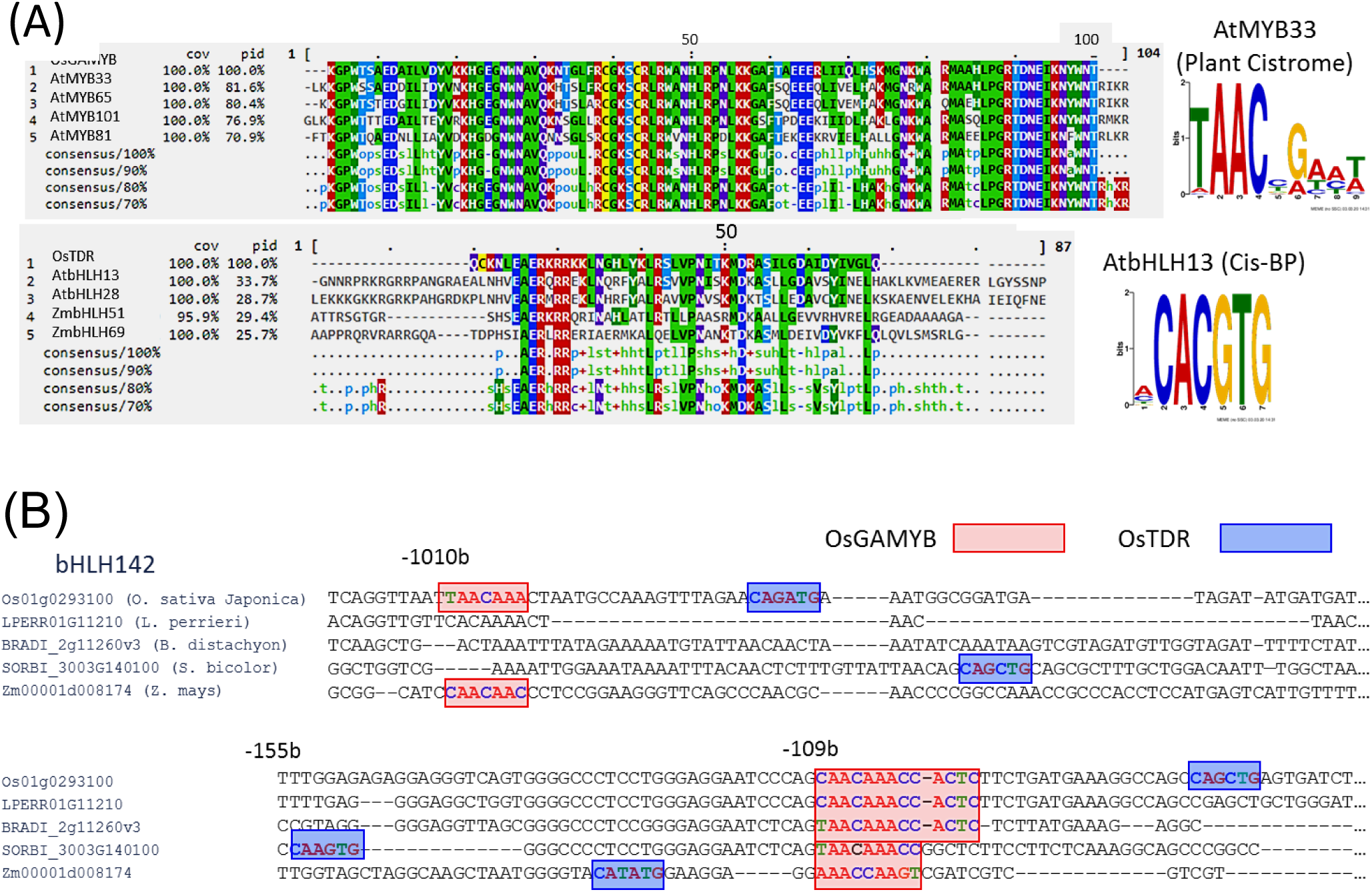
Inferring DNA-binding motifs of *GAMYB* and *TDR*, and identifying their TF binding sites in the promoter sequence of *bHLH142*. (A) DNA-binding domain (DBD) sequences of GAMYB (top) and TDR (bottom) and their top 4 similar DBDs of TFs with known motifs are shown. The logo of the known motif of *AtMYB33* (/*AtbHLH13*) that has the best similar score with *GAMYB* (/*TDR*) is shown at the right panel. The two known motifs (PWMs) are from the Plant Citrome and Cis-BP databases. (B) Alignment of promoter sequences of bHLH142 and four orthologues are shown where two regions in the promoters have been identified as putative TF binding sites of GAMYB and TDR, respectively. The binding site of GAMYB (/TDR) is enclosed by a red (/blue) box.

### *In vivo* and *in vitro* binding assays of GAMYB and TDR to *bHLH142* promoter

Based on our data in Fig 1C and 2, we speculated that GAMYB modulates *bHLH142* transactivation. A transient promoter assay (TPA) indicated that GAMYB activates the expression of *bHLH142pro-LUC* 3.8-fold in the protoplast of rice (Fig. 3A). This experiment was repeated three times to confirm the observation. We then conducted EMSA assays to verify the binding of GAMYB to the core motif of MYB *in vitro*. Because the 1.5 kb promoter of *bHLH142* has two regions consisting of putative GAMYB and TDR binding motifs, respectively (Fig. 2B), two probes were synthesized to perform EMSA assays: probe 1 was a 66-bp sequence from -59 to -124 bp from TSS and probe 2 was a 58-bp sequence from -977 to -1034 bp from TSS. When labeled probe 1 was incubated with GAMYB-MBP protein, a retarded band was observed. In addition, the specificity was confirmed by the competition experiment (Fig. 3B). We also performed an EMSA assay with a competitor, further showing that TDR-MBP protein binds to the E-box close to the MYB box on the promoter of *bHLH142* probe 1 (Fig. 3C). Similarly, when probe 2 was incubated with the GAMYB and TDR protein, respectively, a retarded band was also observed (Fig S3A, B). These data clearly demonstrated that GAMYB and TDR bind to the MYB motif and E-box on the probe 1 and probe 2 of *bHLH142* promoter, respectively.

**Fig. 3.**
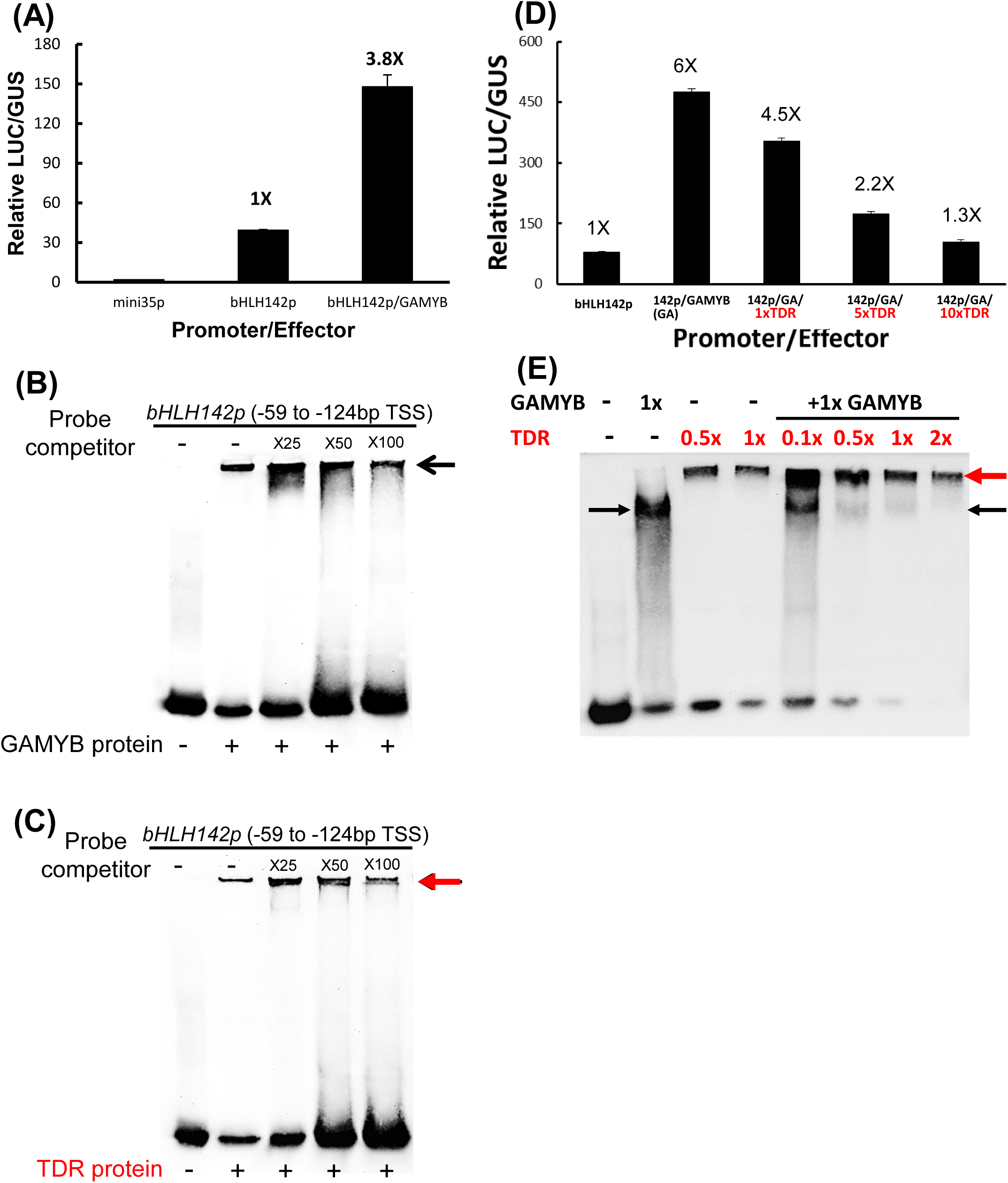
*In vivo* and *in vitro* assays of binding of GAMYB and TDR to *bHLH142* promoter. (A) Transactivation of the *bHLH142* promoter sequence (3 kb) fused to the firefly luciferase gene (*LUC*) reporter gene by GAMYB in rice protoplasts. Different effectors were co-transfected with the reporter and internal control plasmid (pBI221). The data represent means of three independent transient transformations. Transient transformation without the effector plasmid (mini35p) was used as a negative control. Bars indicate the SD of three biological replicates. The experiment was repeated three times. (B) EMSA assay of binding of GAMYB to the promoter region of *bHLH142*. Probe 1 at -59 to -124bp from the transcription start site (TSS) of *bHLH142* promoter was synthesized. (C) EMSA assay of binding of TDR to the promoter of *bHLH142* probe 1. The arrows indicate the protein and DNA complex (B,C). The EMSA experiments were repeated three times. (D) Promoter transient assay indicated that TDR protein inhibited GAMYB modulation on *bHLH142* promoter-*LUC* activity. Different molar ratios of TDR (1x, 5x, and 10x) and GAMYB proteins were used for transient assays. The experiment was repeated three times. Bars indicate the SD of three biological replicates. (E) EMSA data indicated that TDR inhibited GAMYB binding on *bHLH142* promoter probe 1. The black and red arrows indicated GAMYB/ DNA complex and TDR/ DNA complex, respectively.

**Fig. 4.**
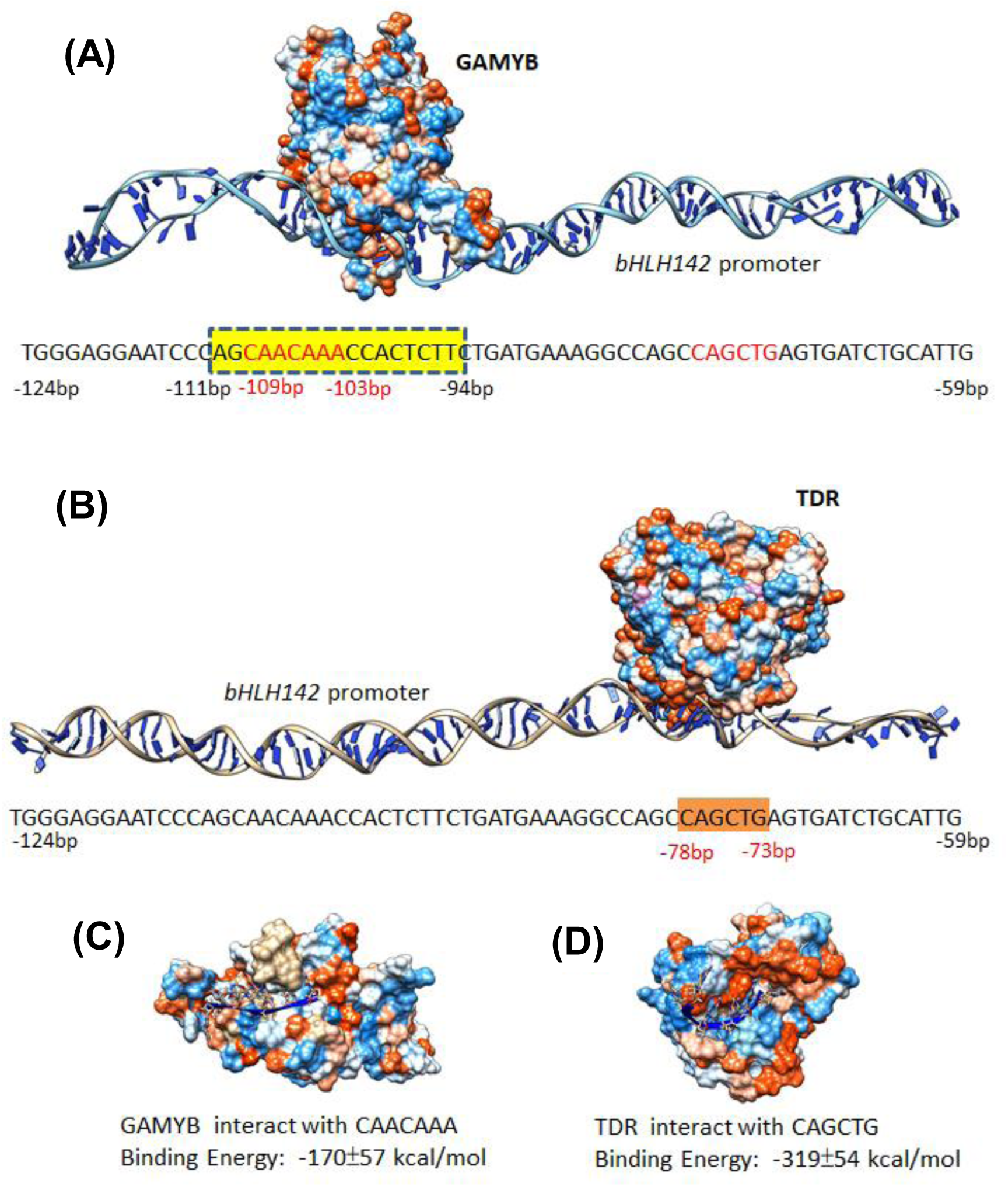
Molecular dynamics simulations of protein/DNA interaction between GAMYB monomer or TDR homodimer and *bHLH142* DNA. (A) Binding of the predicted amino acid patches on GAMYB to the *bHLH142* promoter sequence AGCAACAAACCACTCTTC (−94 to -111 bp from TSS). (B) Binding of the predicted amino acid patches on TDR homodimer to the *bHLH142* promoter sequence CAGCTG (−73 to -78 bp from TSS). (C) Interaction between GAMYB and the CAACAAA motif with a binding energy of -170±57 kcal/mol. (D) Interaction of TDR with an E-box (CAGCTG) with a binding energy of -319±54 kcal/mol.

As a previous study indicated that HvGAMYB regulates hydrolase genes during barley seed development and seed germination in a GA-dependent manner (Gubler *et al*., 1999), we examined whether GAMYB modulation of *bHLH142* expression is also GA_3_ dependent. Our TPA data indicated that adding GA_3_ from 0 to 10 µM did not affect the transactivation of *bHLH142pro*-*LUC* (Fig. S4A). Thus, GAMYB modulates *bHLH142* during pollen development independent of GA_3_. Because bHLH142 acts downstream of GAMYB and UDT1 (Fig. 1B), we were curious whether UDT1 also modulates *bHLH142* promoter activity in the same way as GAMYB. TPA assay showed that UDT1 protein cannot directly transactivate *bHLH142* (Fig. S4B). As TDR is located downstream of the GAMYB regulatory pathway(Liu *et al*., 2010), we were also curious whether GAMYB can modulate *TDR* promoter. Again, our TPA assay indicated that GAMYB does not directly regulate *TDR* (Fig. S4C).

### TDR represses GAMYB-mediated transcriptional activation of *bHLH142*

As transgenic rice overexpressing *TDR* suppressed the expression of *bHLH142* (Fig. 1E, F), we hypothesized that TDR inhibits or interferes with the binding of GAMYB on the promoter of *bHLH142*. To verify this hypothesis, we carried out TPA using different molar ratios of TDR and GAMYB proteins. The results showed that GAMYB increased *bHLH142*pro-*LUC* activity by up to 6-fold in the absence of TDR protein, as expected. However, with addition of 1-, 5- and 10-molar ratios of TDR protein in the assays, the activity of *bHLH142*pro-*LUC* decreased progressively to 4.5-, 2.2- and 1.3-fold in comparison to the control activity (Fig. 3D). We further performed EMSA experiment to show the competition between GAMYB and TDR TFs on the promoter of *bHLH142*. Results with probe 1 indicated that when adding GAMYB protein only produced the protein/DNA complex with lower molecular weight (as indicated by black arrow). Adding TDR only, formed larger protein/DNA complex (as indicated by red arrow) due to the fact that TDR forms dimer for binding to its promoter. Adding 0.1 x TDR protein, showed strong binding of TDR/DNA complex and slightly inhibition of GAMYB binding on *bHLH142*. However, adding >1 x TDR strongly suppressed GAMYB binding on *bHLH142* promoter (Fig. 3E). These data strongly suggest that TDR acts as a repressor by inhibiting GAMYB modulation on *bHLH142*. In short, both *in vivo* and *in vitro* experiments suggest that TDR binds to the *bHLH142* promoter and represses the transactivation of *bHLH142* through GAMYB.

### Molecular dynamic simulation shows binding of GAMYB and TDR proteins to *bHLH142* promoter

To illustrate how GAMYB and TDR interact to regulate the transactivation of *bHLH142* by binding to its promoter, Protein Modeling and Molecular Dynamic (MD) simulation were conducted to determine their protein-DNA interactions. The 3D structures of GAMYB and TDR, were obtained by *ab initio* protein modelling methods through MD simulation in an aqueous environment(Nelson *et al*., 1996). GAMYB, which belongs to the MYB family, is able to bind DNA as a monomer, presumably due to the involvement of an additional N-terminal tail that wraps around the DNA and makes base contacts in the minor groove (Hosoda *et al*., 2002; Ogata *et al*., 1994). In contrast, TDR forms a homodimer in which the two monomers make identical contacts with the DNA. TDR contains a bHLH domain of 281–330 amino acids. The HLH region comprises two amphipathic α-helices separated by a loop of variable length and sequence, allowing the formation of homodimers or heterodimers (Wei and Chen, 2018).

GAMYB and TDR proteins possess electrical and hydrophobic interactive surface patches for protein/DNA interaction. GAMYB protein is predicted to bind to the AGCAACAAACCACTCTTC sequence on the *bHLH142* promoter (−94 to -111 bp from TSS) (Fig. 4A). Our modeling also demonstrated that TDR binds an E-box (CAGCTG) on the *bHLH142* promoter at -73 to -78 bp from TSS (Fig. 4B). Normally, dissociation Gibbs free energy (ΔG) is used to measure the binding affinity. The binding free energies for the binding affinity of GAMYB to CAACAAA and the binding of TDR to CAGCTG on the *bHLH142* promoter were −170 kcal/mol and −319 kcal/mol, respectively (Fig. 4C, D). We found that TDR dimer/DNA complex is more stable than GAMYB/DNA complex. Thus, whenever the GAMYB protein level is high, it will transactivate *bHLH142* gene expression; however, whenever the TDR protein level is high, it acts as a repressor because it has a higher binding affinity for more stable complex formation.

### Spatial expression patterns of *bHLH142*

We used RNA fluorescence *in situ* hybridization (FISH) for direct visualization of the spatiotemporal expression pattern of *bHLH142* (Huang *et al*., 2019; Javelle *et al*., 2011). A strong RNA FISH signal was observed in anthers (Fig. 5A); red strong fluorescence signals of *bHLH142* transcript (as indicated by arrows) were detected in the tapetal layer, middle layer, and meiocytes, with some weak signals detected in the epidermal outer layer and the endothecium of the anther of the WT. In contrast, no apparent *bHLH142* signal was detected in the anther of *ms142*. In the anthers of *udt1* mutant only weak signals of *bHLH142* mRNA, presumably from the GAMYB-dependent pathway, were seen in the tapetal layer but not in the meiocytes. In the anther of *gamyb-2*, some *bHLH142* FISH signals were detected in the meiocytes and in the tapetal cells (Fig. 5A). Taken together, these data suggest that FISH signals in the anther of *gamyb-2* mutant mimic the UDT1-dependent regulatory pathway that regulates *bHLH142* in both tapetal layer (sporophyte) and meiocyte (gametophyte). However, the *udt1* mutant showed the GAMYB-dependent regulation of *bHLH142* expression only in the tapetal layer. Colorimetric detection of the same target genes using Dig-labelled probe and NBT/BCIP detection of the same paraffin blocks of FISH also showed an identical ISH and FISH signal for each mutant as well as the WT (Fig. S5). Differential interference contrast microscopy images of the anther morphology showed that the anther of *udt1* exhibited degenerated meiocytes at the early stage of meiosis (Fig. S6).

**Fig. 5.**
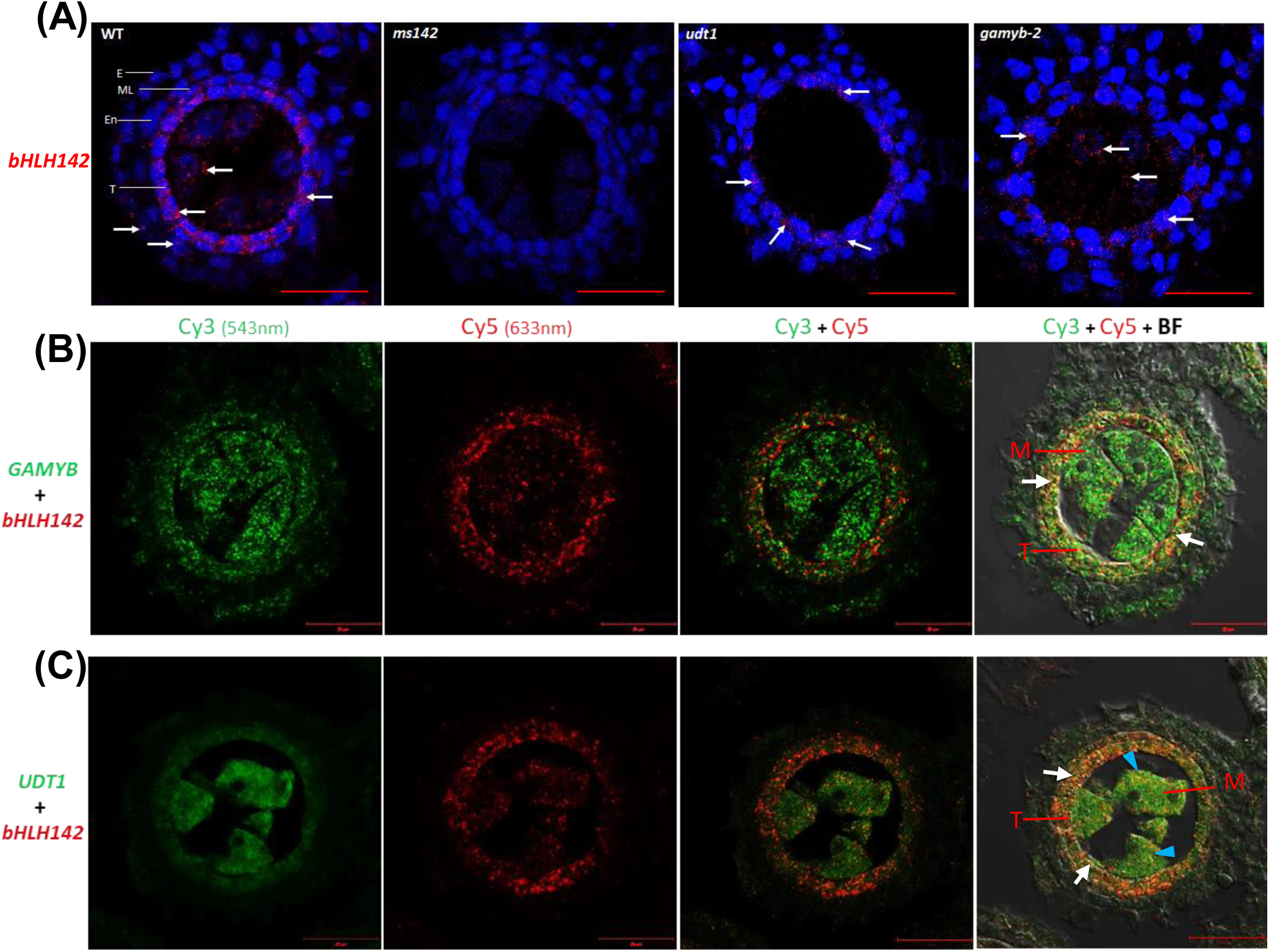
RNA fluorescent *in situ* hybridization (FISH) study of the expression patterns of *bHLH142* in mutants and two-color fluorescent *in situ* hybridization (DISH) study of the co-expression of *UDT1* and *GAMYB* with *bHLH142* in anthers at early meiosis. (A) RNA single color FISH showing the expression pattern of *bHLH142* in the anthers of WT, *ms142, udt1* and *gamyb-2* mutants during early meiosis. Anther tissues were stained with TSA working solution containing Cyanine 5 Plus Amplification Reagent (300x dilution). Ex/Em: 633/670 nm. The arrow indicates the FISH signal (red color). Tissues were counter stained with DAPI (4’,6-diamidino-2-phenylindole). E, epidermis; En, endothecium; ML, middle layer; T, tapetum. Bars = 20 µm. (B) DISH showing coexpression of *GAMYB* and *bHLH142* in the tapetal layer of the anther. Green and red spots show fluorescent signals of *GAMYB* and *bHLH142*, respectively. Yellow spots show the co-expression of *GAMYB* and *bHLH142* mRNAs in the tapetal cells (white arrows). (C) DISH showing co-expression of *UDT1* and *bHLH142* in the tapetal layer and meiocytes of the anther. Green and red spots show fluorescent signals of *UDT1* and *bHLH142*, respectively Yellow spots show co-expression of *UDT1* and *bHLH142* in the tapetal cells (white arrows) and meiocytes (blue arrow heads). Pictures were taken under 100X objective oil immersion lens. T, tapetal layer; M, meiocytes. Bars = 20 µm.

Moreover, our two-color fluorescence *in situ* hybridization (DISH) demonstrated that both *GAMYB* and *bHLH142* genes co-expressed in the same tissue slice of WT anther at the early meiosis stage. Green spots indicate the fluorescence signal of *GAMYB* mRNA expressed in the anther walls and meiocytes (Fig. 5B). Red spots indicate *bHLH142* expressed in the tapetal layer and partly in the meiocytes (Fig. 5B). Co-expression of *GAMYB* and *bHLH142* mRNAs was found only in the tapetal cells (Fig. 5B yellow spots). These data reveal that GAMYB regulates *bHLH142* transcript specifically in the tapetal layer. Dye-swap of the two probes showed similar fluorescence signals (Fig. S7). In addition, we performed DISH to detect the expression of both *UDT1* and *bHLH142* in anther, which suggests the co-expression of these two genes in tapetal cells and meiocytes (Fig. 5C). Taken together, the analyses of RNA FISH using *gamyb-2* and *udt1* mutants and DISH with TNG67 demonstrated that *bHLH142* mRNA is indeed differentially expressed in different tissues. We hypothesize that GAMYB and UDT1 TFs regulate *bHLH142* and might have unique biological functions in these two pollen developmental pathways.

### Expression and putative functions of *bHLH142* in the GAMYB- and UDT1-dependent regulatory pathways

Next, we analyzed the expression of some known pollen marker genes that are significantly downregulated in *ms142* and examined whether their expressions are altered in the anthers of *gamyb-2* and *udt1* mutants, respectively. Interestingly, some marker genes were downregulated in *gamyb-2* and *ms142*, but not significantly altered in *udt1* mutant (named the GAMYB-dependent pathway). On the other hand, some were downregulated in *udt1* and *ms142*, but not significantly changed in *gamyb-2* (named the UDT1-dependent pathway). Our qRT-PCR analysis indicated that the transcripts for some marker genes associated with lipid transport, such as *YY1, OsC4*, and *LTP2*, were sharply decreased in the anthers of *ms142* and *gamyb-2* mutants but not in that of *udt1* (Fig. 6A). However, *RAD51*, a marker gene involved in meiotic homologous recombination (Shinohara *et al*., 1992), was downregulated in both *udt1* and *ms142*, but not in *gamyb-2* (Fig. 6B). Also, *PTC1* and *EAT1* were greatly downregulated in *udt1* and *ms142*, but not in *gamyb-2* (Fig. 6B). The expression of some other genes, such as *TDR, MYB80*, sporopollenin biosynthesis genes (*CYP703A3* and *CYP704B2*), and *PKS1* (*YY2*), were simultaneously downregulated in these three mutants (Fig. 6C). Collectively, these results reveal that some known pollen markers acting downstream of bHLH142 are dependent on the UDT1- or GAMYB-pathway, while some others are dependent on both pathways during rice pollen development.

**Fig. 6.**
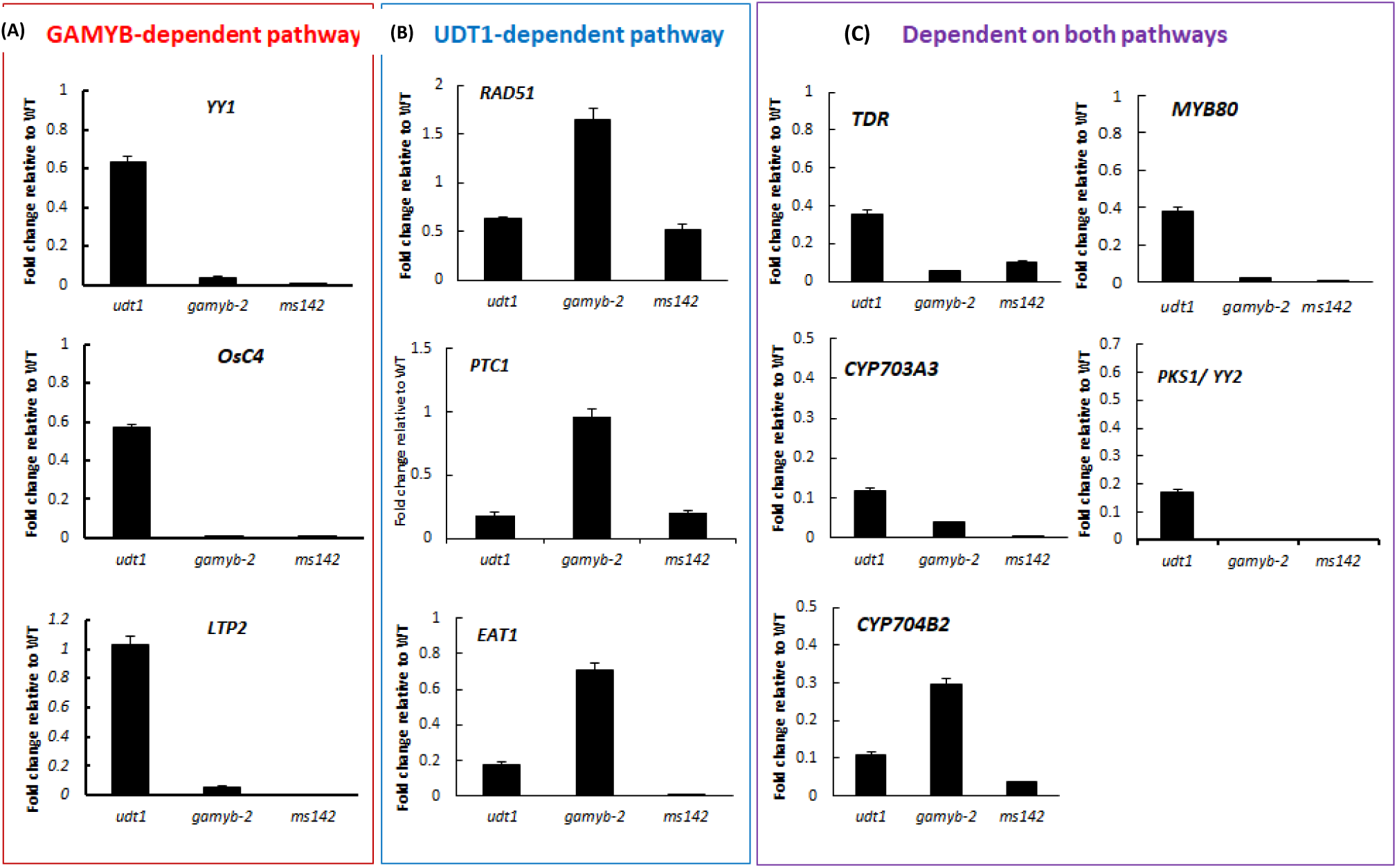
Dependence of the expression of pollen marker genes on GAMYB, UDT1, or both pathways. Expression patterns of some pollen marker genes in the anthers of *udt1, gamyb-2*, and *ms142* mutant lines were studied using qRT-PCR. (A) *YY1, OsC4*, and *LTP2* were found dependent on GAMYB- and bHLH142-regulatory pathways. (B) *RAD51, PTC1*, and *EAT1* were dependent on UDT1- and bHLH142-regulatory pathways. (C) *TDR, CYP703A3, Cyp704B2, MYB80*, and *PKS1/YY2* were dependent on both pathways. Gene expression levels were normalized to *Ubi5*, and the fold change relative to the respective Wild-type is presented. Error bars indicate the SD of three technical replicates. Each experiment was repeated three times.

## Discussion

GAMYB, UDT1, bHLH142, and TDR are master transcriptional regulators of rice tapetal and pollen development at the meiosis and early microspore stages. Previous studies suggest that GAMYB and UDT1 work in two parallel pathways (Liu *et al*., 2010) and bHLH142 acts downstream of GAMYB and UDT1 (Ko *et al*., 2014). This study further demonstrates that bHLH142 acts as a “hub” of these two pathways. We showed that GAMYB directly modulates *bHLH142* transcription by binding to its promoter (Fig. 3A, B), but UDT1 does not directly transactivate *bHLH142* expression (Fig. S4B). Presumably, UDT1 (bHLH164) needs to interact with another bHLH to form a heterodimer to co-modulate *bHLH142* expression. This is similar to our previous discovery that two bHLHs, bHLH142 and TDR (bHLH5), form a heterodimer to co-modulate another downstream bHLH gene, *EAT1* (*bHLH141*) (Ko *et al*., 2014). Alternatively, there is an unknown TF acting downstream of UDT1 that might directly modulate *bHLH142*. In Arabidopsis, the two TFs MS188 (MYB80) and AMS (a bHLH, TDR homolog in rice) form a complex to activate the expression of the fatty acid hydrolase gene *CYP703A3* for sporopollenin biosynthesis (Xiong *et al*., 2016). Clearly, spatial and temporal coordination among the regulatory TFs is essential for rice pollen maturation. Using mutagenesis, FISH and DISH, we demonstrated that GAMYB specifically modulates *bHLH142* expression in the tapetal cells (Fig. 5). Our qRT-PCR data showed that *YY1* (*LTP45*), *LTP2*, and *OsC4 (LTP44)* transcripts are downregulated in the anther of *gamyb-2* and *ms142*, but not significantly altered in *udt1* (Fig. 6A). These results suggest that the expression of these lipid transfer genes is dependent on bHLH142 via a GAMYB-dependent pathway. UDT1 modulates *bHLH142* expression in the tapetal cells and meiocytes (Fig. 5), indicating that the UDT1-dependent pathway regulates *bHLH142* for sporophytic and gametophytic development during rice pollen development (Fig. 5). *RAD51* is downregulated in both *udt1* and *ms142* mutants (Fig. 6B) and might be associated with meiosis defective in *udt1* and *ms142* mutants. Two TFs, *PTC1* and *EAT1*, are involved in UDT1-dependent pathway (Fig. 6B). In other words, the UDT1-dependent pathway indirectly modulates *bHLH142* in broad tissue compartments including sporophyte (tapetum) and gametophyte (meiocyte). This result is in agreement with the function of bHLH142 in tapetal layer differentiation(Fu *et al*., 2014) and its regulation of various metabolic pathways during rice pollen development (Ranjan *et al*., 2017).

This study also clearly demonstrated that TDR plays an important role in regulating the homeostatic gene expression of *bHLH142*. Transgenic lines overexpressing *TDR* showed a differential *TDR* expression pattern and the *TDR* and *bHLH142* transcript levels were negatively correlated (Fig. 1F, G). Our TPA demonstrated that a high amount of TDR protein can repress the transactivation of *bHLH142* expression by GAMYB (Fig. 3D). Moreover, EMSA assay suggests that TDR directly binds to the E-box adjacent to a MYB motif on the *bHLH142* promoter region (Fig. 3C), which in turn suppressed the transactivation of *bHLH142* expression by GAMYB in a competitive manner (Fig. 3E). Hence, based on this and previous studies, we propose that TDR has at least three molecular functions during rice pollen development. First, it controls homeostasis of *bHLH142* by inhibiting the transactivation of *bHLH142* by GAMYB (Fig. 3). Second, TDR interacts with bHLH142 to form a heterodimer (bHLH142/TDR) to co-modulate *EAT1* expression (Ko *et al*., 2014), and a sharp upregulation of *EAT1* mRNA is found at the YM stage (Fig. 1A). Moreover, overexpressing *bHLH14*2 caused premature upregulation of *EAT1*(Ko *et al*., 2017). Third, TDR interacts with EAT1 (Ji *et al*., 2013; Ko *et al*., 2014; Niu *et al*., 2013) in regulating the expression of the downstream genes for pollen maturation.

Based on the results from this and others studies, we propose an updated regulatory model of rice pollen development (Fig. 7). First, bHLH142 acts downstream of UDT1 and GAMYB and works as a “hub” in these two pollen development pathways (Fig. 1B). Second, GAMYB, but not UDT1, can directly modulate the transactivation of *bHLH142* promoter (Fig. 3A, Fig. S2B). Third, GAMYB modulates *bHLH142* at the early stage of meiosis (Fig. 1A). Consistently, from the late meiosis to early YM stages, a high level of *bHLH142* transcript accumulates in the cytosol of tapetum cells (Ko *et al*., 2014). Fourth, in the meantime, TDR regulates the homeostasis of *bHLH142* expression by binding to an E-box adjacent to the MYB motif on its promoter region and inhibits GAMYB’s modulation of *bHLH142* (Fig. 3). At the YM stage, TDR interacts with bHLH142 to co-modulate *EAT1* promoter as reported previously (Ko *et al*., 2014). EAT1 in turn regulates *AP37* to turn on tapetal PCD (Niu *et al*., 2013) at the YM stage. Finally, both FISH and DISH data reveal the modulation of *bHLH142* by GAMYB in the tapetal layer (Fig. 5) and the GAMYB-dependent pathway functions in lipid transport (Fig. 6A). However, the UDT1-dependent pathway plays more important roles in meiosis and tapetal programmed cell death (Fig. 6B). Overall, the present study underlines the importance of a precise and coordinated expression of these regulatory TFs spatially and temporally for pollen development.

**Fig. 7.**
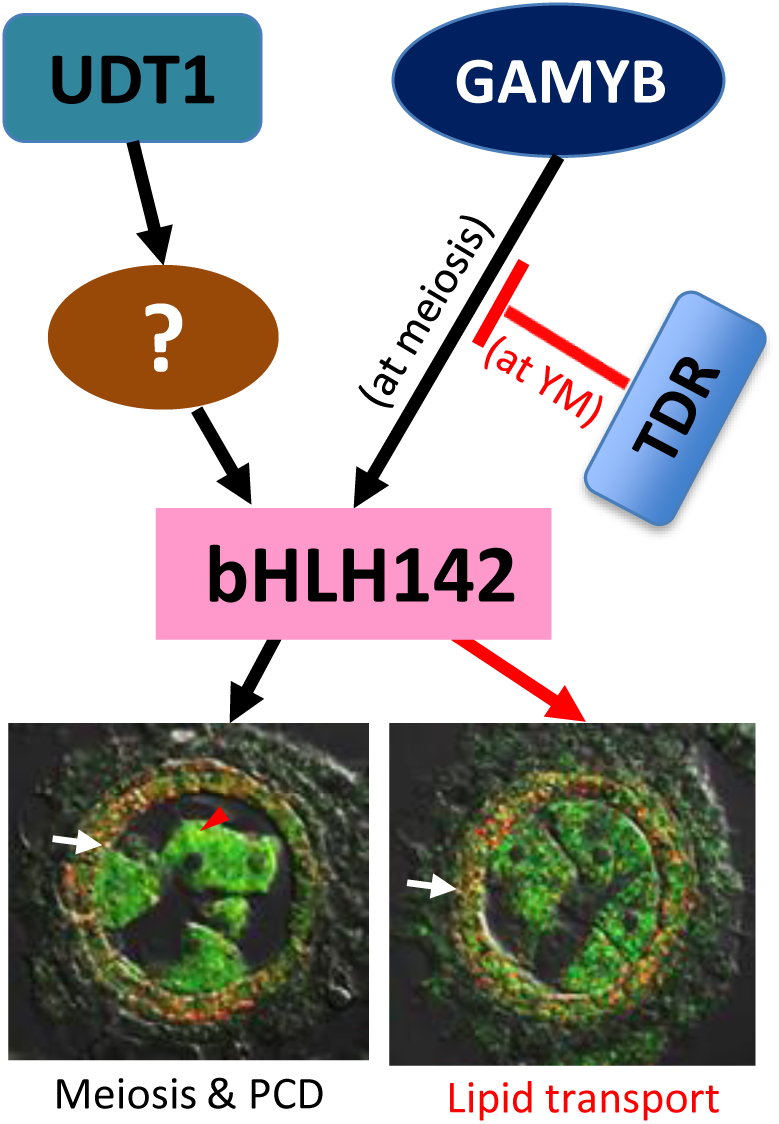
Model of this study. TF of bHLH142 acts downstream of UDT1 and GAMYB and works as a “hub” in these two pollen pathways. GAMYB can directly modulate the transactivation of the *bHLH142* promoter, but UDT1 cannot. GAMYB modulates *bHLH142* at the early stage of meiosis, but the modulation is repressed by TDR at the young microspore stage to keep the homeostasis of *bHLH142* gene expression to ensure normal pollen development. Our FISH and DISH data reveals that GAMYB modulates *bHLH142* in the tapetal layer and GAMYB-dependent pathway functions in lipid transport. However, the UDT1-dependent pathway regulates *bHLH142* in the tapetal layer and meiocytes and plays several other roles in meiosis and tapetal programmed cell death. YM, young microspore stage; M, meiocytes; T, tapetum; PCD, programmed cell death.

### Accession numbers

Sequence data from this article can be found in the GenBank/EMBL database under the following accession numbers: *bHLH142* (Os01g0293100), protein (NP_001042795.1); *GAMYB* (Os01g0812000); *UDT1* (Os07g0549600), and *TDR* (Os02g0120500). Additional loci are presented in Table S1.

## Acknowledgements

We would like to thank the TRIM, Tos17, and Postech mutant libraries for providing mutant seeds for this study. We thank the Academia Sinica Transgenic Plant Laboratory for transforming the OE-*TDR* rice. We thank the GMO greenhouse of the Biotechnology Center in Southern Taiwan (AS-BCST) for providing space and technical support. We thank the Confocal Microscope Core Facility of AS-BCST for confocal microscopy and imaging assistance. We thank the DNA Sequencing Core Facility of the Institute of Biomedical Sciences of Academia Sinica for providing DNA sequencing services. We appreciate Miranda Loney for English editing. This work was supported by grants of AS-BCST and partly by The Ministry of Science and Technology to S.S. Ko (MOST 107-2313-B-001-005), and AS-109-TP-10, Academia Sinica to W.H. Li.

## Author contributions

S.-S.K. conceived and designed the research. M.-J.L. discovered *bHLH142* promoter is directly modulated by GAMYB. Y.-C. H. and C.-P. Y. performed MD simulation and TFBS analysis. M.-J.L., T.-T.Y., Y.-J.L., H.-C.H., T.-K.C., and C.-M.J. carried out the experiments. S.-S.K., M.-J.L., Y.-C. H., C.-P. Y. analyzed and prepared the data. M.S.-B.K. and W.-H.L. suggested on the experiments. S.-S.K., M.S.-B.K., Y.-C. H., and W.-H.L. wrote the article.

## Supplementary data

Table S1. Primers used in this study.

**Fig. S1**. Phenotype of the TNG67 wild-type and overexpressing *GAMYB* transgenic line. (A) Construction map of overexpressing *GAMYB* driven by *Ubiquitin* promoter. (B) Genomic PCR confirmed insertion of Ubi::GAMYB in transgenic lines. (C) Plant phenotype at the seed maturation stage. (D) Spikelet at one day before anthesis. (E) Pollen stained with 1% I_2_/KI solution. Left panel is WT, right panel is OE-*GAMYB* #6. Scale bars, 20 cm (C), 1mm (D), 50 um (E).

**Fig. S2**. Gene expression level of *GAMYB* and *UDT1* were not altered in the anther of *OE-TDR* transgenic lines. RNA of anthers at the meiosis stage was collected from the Wild-type (TNG67) and 15 independent *OE-TDR* transgenic lines. The Pearson correlation coefficient between *GAMYB* and *UDT1* mRNA is -0.27. Bars indicate SD of mean from three technical replicates. n = 16 lines.

**Fig. S3**. EMSA results indicated GAMYB and TDR TFs bind to *bHLH142* promoter sequences at -977 to -1034 bp from TSS. (A) GAMYB protein bound to the promoter region of *bHLH142*. (B) TDR protein bound to the promoter region of *bHLH142*. Probe 2 at -977 to -1034 bp from the transcription start site (TSS) of *bHLH142* promoter was synthesized. Arrow indicated protein and DNA complex. The experiment was repeated three times.

**Fig. S4**. Transient promoter analysis. (A) GAMYB modulation of *bHLH142* promoter is independent of GA_3_. (B) UDT1 does not directly modulated by *bHLH142*pro-*LUC* activity. c, *TDR* promoter does not modulated by GAMYB TF. This experiment was repeated three times.

**Fig. S5**. Traditional Dig-labelled antisense probe of *bHLH142* in the anthers of WT, *ms142, udt1* and *gamyb-2* mutants at early meiosis stage. Transverse sections of anther were hybridized with antisense or sense Dig-labeled probe of *bHLH142*. Bars = 20 µm.

**Fig. S6**. DIC of the anthers of WT, *ms142, udt1* and *gamyb-2* mutants at the early meiosis stage for RNA FISH analyze *bHLH142* mRNA expression patterns. The arrows indicates degenerated meiocytes in *udt1* mutant. Bars = 20 µm.

**Fig. S7**. Two-color fluorescent ISH (DISH) of *GAMYB* and *bHLH142* mRNAs in the anther of TNG67 at the early stage of meiosis. DISH experiment performed reciprocal staining of TSA Plus Cy3 and Cy5 to show RNA fluorescence signals. Fluorescence signals were observed under a Zeiss LSM710 confocal microscope equipped with a T-PMT, using Cy3 or Cy5 filter at excitation/emission wavelengths of 543/600 nm and 633/670 nm, respectively. Pictures were taken under 40X objective lens. Bars = 20 µm.

